# Mechanotransduction-Aware Causal Omics on Tissue Scaffolds: A Controlled Mechanochemical Framework for Identifying Disease Genes Beyond Pure Omics Analysis

**DOI:** 10.64898/2026.04.19.719528

**Authors:** Tao Xu, Zhixin Hu, Xiaodian Sun, Momiao Xiong

## Abstract

Omics-based disease-gene discovery is typically performed as if molecular states evolve independently of tissue mechanics. Most current pipelines analyze transcriptomic or multimodal molecular data alone and identify abnormal genes using differential expressions, latent trajectories, or association-based recovery under treatment. However, in mechanically active tissues, gene expression is shaped not only by internal regulatory networks but also by mechanotransduction arising from strain, curvature, force transmission, and scaffold geometry. This raises a fundamental question: should disease-gene identification in tissues be treated as a pure omics association problem, or as a causal mechanochemical inference problem?

We introduce a mechanotransduction-aware causal omics framework on a Cosserat tissue scaffold. Gene expression evolves through intrinsic regulatory dynamics, spatial diffusion, external control, and a mechanotransduction term driven by scaffold mechanics. To distinguish causation from association, we define a hidden mechano-drug rescue channel in the true data-generating system and compare predictive models that either include or omit mechanotransduction. We show that association-based rankings can incorrectly elevate downstream homeostatic or repair genes, even when the disease gene is the true direct mechanochemical target. By contrast, a causal ranking based on reconstruction of the direct mechanotransduction intervention effect correctly identifies the disease gene as the strongest beneficiary.

These results argue that popular pure-omics analysis is insufficient for disease-gene discovery in mechanically structured tissues. Mechanotransduction should be modeled as part of the causal structure of tissue biology rather than treated as a secondary covariate or omitted entirely.

## 1. Introduction

Omics analysis has become a standard approach for identifying disease genes, pathways, and treatment response. Differential expression analysis, low-dimensional embeddings, trajectory inference, RNA velocity, and multimodal latent-variable models all attempt to explain biological state transitions from molecular measurements. In most of these frameworks, tissue mechanics is either ignored or treated only indirectly through secondary covariates such as morphology or position.

This omission is problematic in mechanically active tissues (Hallou et al. 2026; Mathavan et al. 2025; Xie et al. 2026; Discher et al. 2005; Engler et al. 2006; Humphrey et al. 2014). Cells in epithelia, luminal surfaces, vessels, organoids, and engineered scaffolds experience curvature, deformation, pressure, force transmission, and elastic energy storage. These mechanical signals are known to influence transcription through mechanotransduction pathways such as YAP/TAZ, integrin–FAK, β-catenin, and TGF-β/Smad (Long et al. 2020; Miranda et al. 2017; Heng et al. 2021; Ríos-López et al. 2023). Consequently, molecular states are not governed solely by gene regulation and biochemical signaling. They are also shaped by tissue mechanics (Editorial 2026; Yang et al. 2026; Cha and Thibeault 2025). This observation changes the nature of disease-gene identification. In a purely molecular view, the task is to find genes most strongly associated with abnormality or recovery. In a mechanochemical view, the task is to identify genes that are causally modulated by mechanically mediated intervention channels. These are not equivalent (Yeo and Selvarajoo 2026; Xiong 2022). A downstream repair gene may exhibit broad changes and dominate association-based rankings, while the true disease gene may be the direct target of a hidden mechanochemical rescue mechanism.

In this work we develop a mechanotransduction-aware causal omics framework that makes this distinction explicit (Feng et al. 2026; Kriel et al. 2026; Guo et al. 2025). We model gene expression on a mechanically structured tissue scaffold using a controlled mechanochemical reaction–diffusion system (Krause et al. 2023; Diez et al. 2024). The true data-generating system includes a hidden mechano-drug rescue channel (Stewart and Gao 2026; Zielinski et al. 2026), while competing predictors either include or omit mechanotransduction. This allows us to separate the mechanochemical truth from the model class used to explain it (Francis et al. 2026). A second contribution of this work is conceptual. We show that ranking genes by total recovery, endpoint fit, or overall trajectory error can fail to identify the true disease gene, even when that gene is the direct mechanotransduction target. Downstream repair or homeostatic genes may exhibit broader or smoother responses and dominate association-style rankings (Song et al. 2025). To address this, we formulate disease-gene identification as a causal intervention-effect reconstruction problem (Wen et al. 2013; Pöyhönen et al. 2025). In our strongest benchmark, the true mechanochemical system contains a hidden mechano-drug rescue channel acting directly on the disease gene. Predictors are then evaluated on their ability to reconstruct the induced causal effect.

We use a controlled computational benchmark (Sawarni et al. 2026) to establish three main points. First, mechanotransduction-aware predictors (Gu et al. 2026) can outperform pure omics predictors on disease-gene prediction under treatment. Second, association-based rankings can misidentify downstream genes as primary beneficiaries. Third, a causal ranking based on reconstruction of direct mechanochemical intervention effects can correctly recover the disease gene as the top beneficiary (Saitow et al. 2026). Taken together, these results support a shift from pure omics analysis toward mechanotransduction-aware causal omics in structured living tissues.

## 2. Methods

### 2.1 Tissue scaffold model

We represent the tissue as a one-dimensional mechanically structured scaffold parameterized by *s* ∈ [0, *L*] (Figure 1).This model treats biological tissue like a **flexible, 1D structural beam** rather than a complex 3D blob. The scaffold can be interpreted as a reduced representation of a curved lumen, epithelial axis, vessel, or engineered tissue support. Its mechanical state (the shape (deformation))is summarized by translational strain *v*(*s, t*),which measures how much the tissue is being stretched or compressed at that point, rotational strain or curvature-twist *u*(*s, t*) measuring how much the tissue is bending or twisting, internal force *N*(*s, t*) that measures the “tension” pulling along the tissue, and internal moment *M*(*s, t*), that measures the “torque” or rotational force inside the tissue. In the present benchmark these fields are prescribed, allowing the inferential role of mechanotransduction to be isolated from the inverse mechanics problem. In this model, biological sensing (mechanotransduction) is how cells on that 1D scaffold “read” the physical signals you just defined (*v, u, N, M*) and turn them into a biological response, like growing, moving, or changing shape.

**Figure 1.**
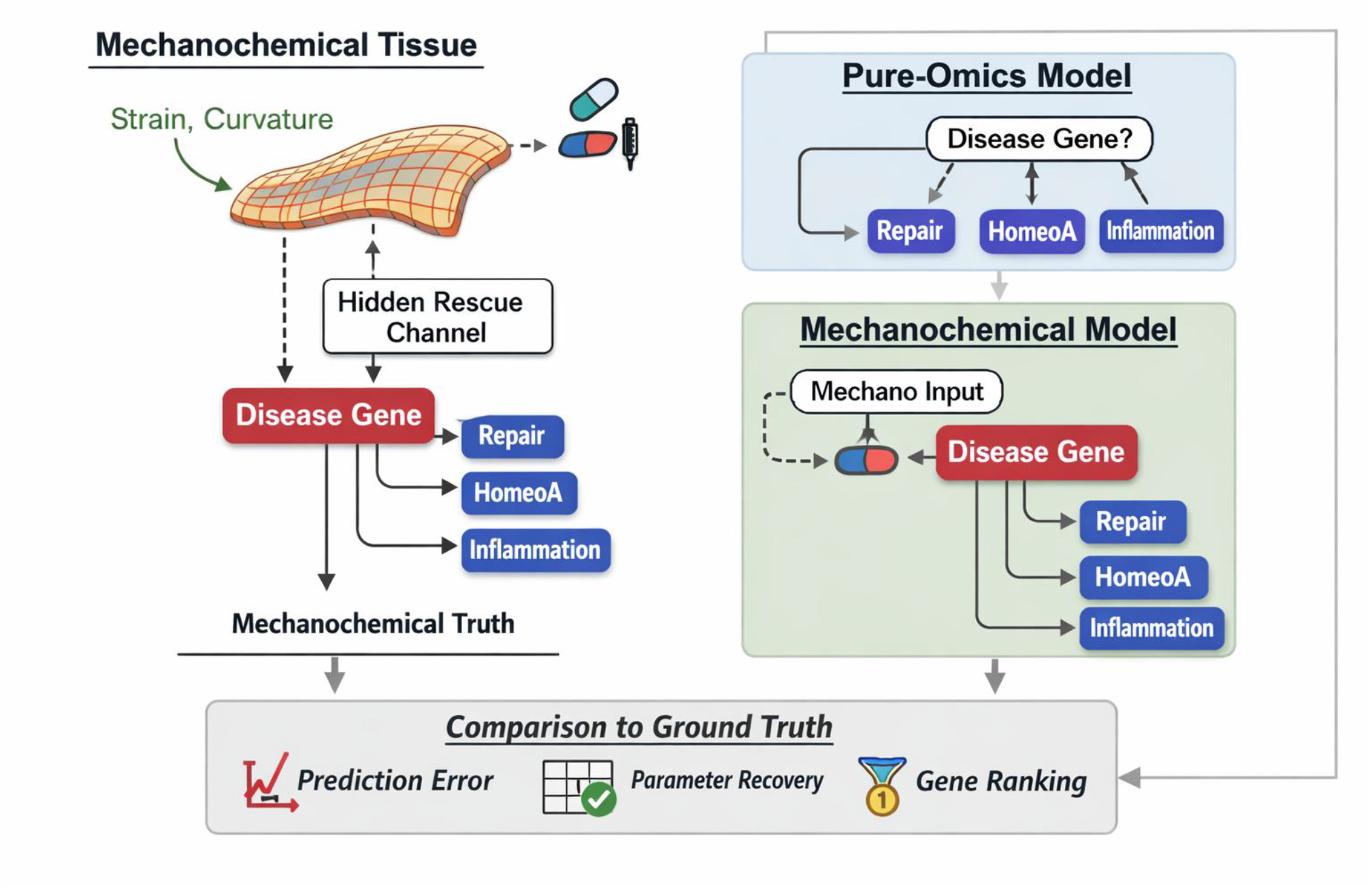
Mechanotransduction-aware causal omics framework. A tissue scaffold carries mechanical fields such as strain and curvature. These fields modulate a hidden drug-conditioned rescue channel acting directly on the disease gene. Predictors with and without mechanotransduction are compared against the same mechanochemical truth.

**Figure 2.**
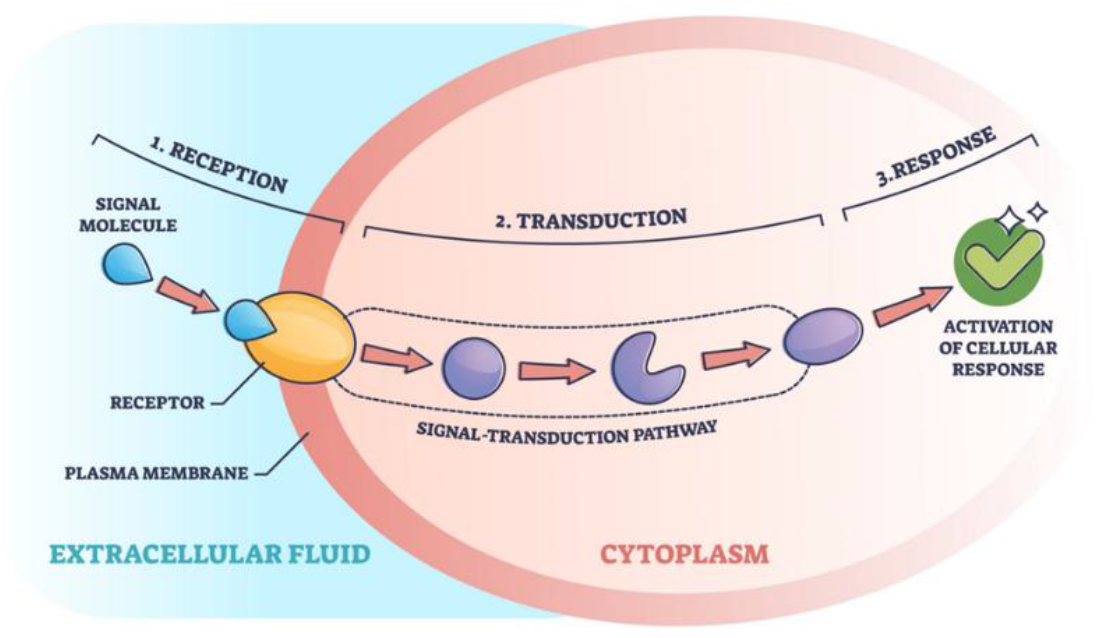
Cell signaling.

Since the physical state is “prescribed” (given to you), the sensing part focuses on two main questions:

1. **What are the cells actually feeling?** The model assumes the cells are sensitive to two specific inputs: **Strain Sensing (*v* and *u*):** Cells feel how much the scaffold is stretched or bent. Think of stretch-activated ion channels in the cell membrane—when the “floor” (scaffold) pulls apart, the channels physically pull open, letting signals in. **Stress Sensing (*N* and *M*):** Cells feel the internal tension or “tug” within the tissue. This is often sensed at focal adhesions, the “hooks” cells use to grab onto their environment.
2. **How do they process that info?** The “inferential role” mentioned in the text suggests the cells are trying to estimate something about their environment based on these signals. For example: **The Set-Point:** Cells often try to maintain a “homeostatic” level of tension. If the sensed *N* (force) is too high, the sensing model might trigger the cell to release enzymes that soften the tissue. **Signal Integration:** The model likely combines the stretching (*v*) and the bending (*u* ) to help the cell “know” if it is part of a straight vessel or a tight curve, which might change how it behaves (e.g., cells on a curve might divide faster to thicken the wall).

### 2.2 Omics state

Let

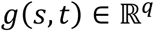

denote the vector of gene expression levels or low-dimensional omics factors. One component is designated as the disease gene, and the remaining components represent non-disease programs such as homeostasis, stress response, repair, and inflammation.

### 2.3 Controlled mechanochemical reaction–diffusion dynamics

The true system is modeled as

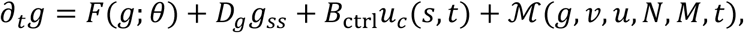

where *F*(*g*;*θ*)is the intrinsic gene regulatory network drift, *D*_*g*_*g*_*ss*_is spatial diffusion or communication, *u*_*c*_(*s, t*)is the external control input, and ℳis the mechanotransduction term. The control field *u*_*c*_(*s, t*)may represent drug application, biochemical stimulation, or optogenetic intervention. The diffusion matrix *D*_*g*_ ⪰ 0models effective spatial communication, such as morphogen spread or gap-junction-mediated smoothing.

This equation describes how a biological system (like a group of cells or a tissue) changes over time and space, influenced by its internal “programming,” its neighbors, and outside forces. Now we explain the equation one component by one component.

- **∂**_*t*_***g* (The Change):** This is the “output.” it represents how the concentration or state of genes (*g*) is evolving at a specific moment in time (*t*).
- ***F***(***g***;***θ***)**(The Internal Logic):** This is the Gene Regulatory Network. It’s the cell’s internal “circuitry”—how genes naturally turn each other on or off based on their own rules (*θ*).
- ***D***_***g***_***g***_***ss***_ **(The Neighborhood):** This is Diffusion. It models how signals spread to nearby cells (like a drop of ink in water). This allows cells to coordinate their behavior rather than acting in total isolation.
- ***B***_**ctrl**_***u***_***c***_(s, t) (**The Remote Control):** This is the Human Intervention. *u*_*c*_ is an external signal—like a drug, a chemical, or a laser (optogenetics)—that we use to force the system to behave a certain way. *B*_ctrl_ determines how sensitive the genes are to that specific input.
- **ℳ**(***g, v, u, N, M***, *t*) **(The Physical Push):** This is Mechanotransduction. It accounts for the “mechanical” side of biology—how physical forces like stretching, squeezing, or stiffness in the environment actually change gene expression.

In summary, imagine you are trying to model how a healing wound closes. The equation says the healing process is a mix of:

What the cells are programmed to do (Internal Logic).

How they talk to the cells next to them (Diffusion).

The medicine you apply to the wound (Control).

The physical tension of the skin pulling on the cells (Mechanical Forces).

### 2.4 Mechanical features and hidden rescue gate

Mechanotransduction depends on local mechanical features. A representative energy-like feature is

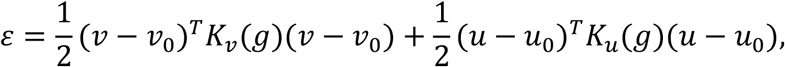

where *K*_*v*_(*g*)and *K*_*u*_(*g*)are gene-dependent constitutive matrices.

The formula describes the mechanical energy (*ε*) stored in a cellular system when it is deformed or moved away from its “rest” state. It models mechanotransduction—the process by which cells convert physical forces into biochemical signals—as a mathematical system where the “cost” of movement depends on the cell’s genetic makeup (Figuren1).

1. **Breaking down the components** The equation is a sum of two quadratic forms, representing different types of mechanical energy (likely related to velocity/viscosity and displacement/elasticity):
  • **State Variables (*v* and *u*):**
    ○ *v* represents the current velocity or rate of change.
    ○ *u* represents the current displacement or deformation.
    ○ *u*_0_ and *v*_0_ are the equilibrium states (where the cell feels “no stress”).
  • **The Deviations (*v*** − ***v***_0_ **and *u*** − ***u***_0_): These terms measure how far the local mechanical features are from their preferred rest state. The larger the deviation, the higher the energy.
2. **The constitutive matrices (*K***_***v***_ **and *K***_***u***_) The terms *K*_*v*_(*g*) and *K*_*u*_(*g*) are stiffness or damping matrices. In a standard physics context, these would be constants. However, in this biological model:
  • **Gene-Dependency (*g*):** The matrices are functions of the expressed genes (g). This means a cell’s physical “stiffness” or “sensitivity” to force is dictated by which proteins (like actin, myosin, or integrins) it is currently producing.
  • **Constitutive Properties:** These matrices define the “material laws” of the cell. They dictate how much energy is generated for a given amount of physical stretch or movement.
3. **Biological interpretation** The equation *ε* represents the local mechanical signal that a cell “senses.”
  • **Energy as a Trigger:** When *ε* increases (because the environment is pushing or pulling the cell way from *u*_0_), it triggers chemical changes.
  • **Feedback Loop:** Because the matrices depend on *g*, the cell can “tune” its sensitivity. If a cell needs to become more rigid to resist flow, it changes its gene expression (*g*), which alters *K*, thereby changing how it perceives the next mechanical force.

In the strongest benchmark, the hidden mechanochemical rescue channel acts directly only on the disease gene:

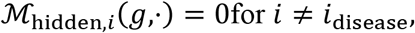

and

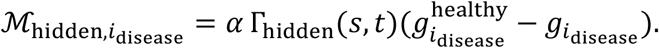

The hidden gate is localized in both space and treatment time:

The hidden gate is localized in both space and treatment time:

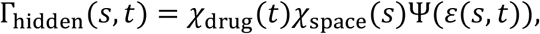

Where χ_space_(*s*) indicates that the drug or effect only happens in a specific part of the body or cell (like a tumor site or a specific joint), treatment time χ_drug_(*t*) indicates that the gate only opens when the drug is actually present or at a specific stage of the disease, Ψis a nonlinear thresholding function. This creates a direct disease-specific rescue pathway that cannot be reconstructed by a purely omics-based predictor unless mechanotransduction is explicitly modeled.

### 2.5 True system versus predictive models

A central principle of the benchmark is the separation of truth from prediction. The **true** data-generating system always includes mechanotransduction. We generate two matched truth trajectories:

- 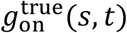: hidden mechanochemical gate enabled
- 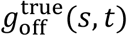: hidden mechanochemical gate disabled

Their difference defines the true intervention effect:

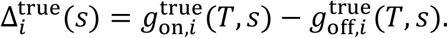

Now we explain this definition. In biological terms, the true intervention effect (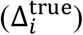) measures the absolute benefit of a treatment that can only be activated by physical force (mechanical stress).

Think of it as a “Clinical Trial in a Computer” that compares two identical scenarios to see if the mechanical “gate” actually works.

1. **The Two Parallel Realities** The model creates two “twins” (trajectories) to see what happens over time:
  - 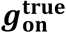 **(The Treated Group):** The patient receives the drug, and the physical forces in their body (like blood pressure or muscle tension) successfully trigger the “rescue gate” to fix the gene.
  - 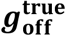 **(The Control Group):** Everything is the same, but the “rescue gate” is intentionally disabled or fails to trigger. The gene expression follows its diseased path.
2. **Real-world Example** In real-world diseases, this “hidden gate” corresponds to mechanosensitive ion channels— specialized proteins on cell membranes that physically open and close in response to mechanical force. **Osteoporosis: The “Use It or Lose It” Gate (Chen et al. 2025)** In bone health, the Piezo1 channel acts as a primary mechanical gate.
  - **The Physical Force:** When you exercise or “pound the stairs,” fluid flows through tiny tunnels in your bones (lacuno-canalicular system), creating fluid shear stress.
  - **The Gate Action:** This physical pressure stretches the cell membrane, physically pulling the Piezo1 gate open.
  - **The Rescue (Benefit):** Once open, it allows calcium to rush into bone cells, triggering genes like Runx2 and Wnt, which tell the body to build more bone and stop breaking it down.
  - **In Disease:** In osteoporosis, this gate often becomes “stiff” or is under-activated due to a lack of physical impact (like bed rest or microgravity), leading to bone loss because the “rescue” signal is never triggered.

A standard drug that treats these diseases chemically might only work 50% as well if the patient is sedentary. In your model’s terms, the “True Intervention Effect” would show that the real cure only happens when the drug is combined with the correct mechanical “push” to open these biological gates.

Next we compare two predictors.

**Mechanotransduction-aware predictor**

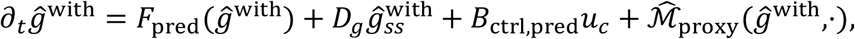

**Pure-omics predictor**

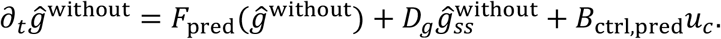

The former has access only to an imperfect proxy of the hidden gate. The latter omits mechanotransduction entirely.

### 2.6 Numerical scheme

The scaffold coordinate is discretized using centered finite differences with Neumann boundary conditions:

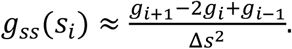

Time stepping is performed by explicit Euler:

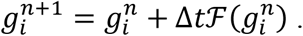

This yields a transparent computational benchmark with known ground truth.

### 2.7 Causal ranking

Association-style rankings based on total trajectory error or total recovery were found to misidentify downstream genes. We therefore define a causal score (Figure 3).

**Figure 3.**
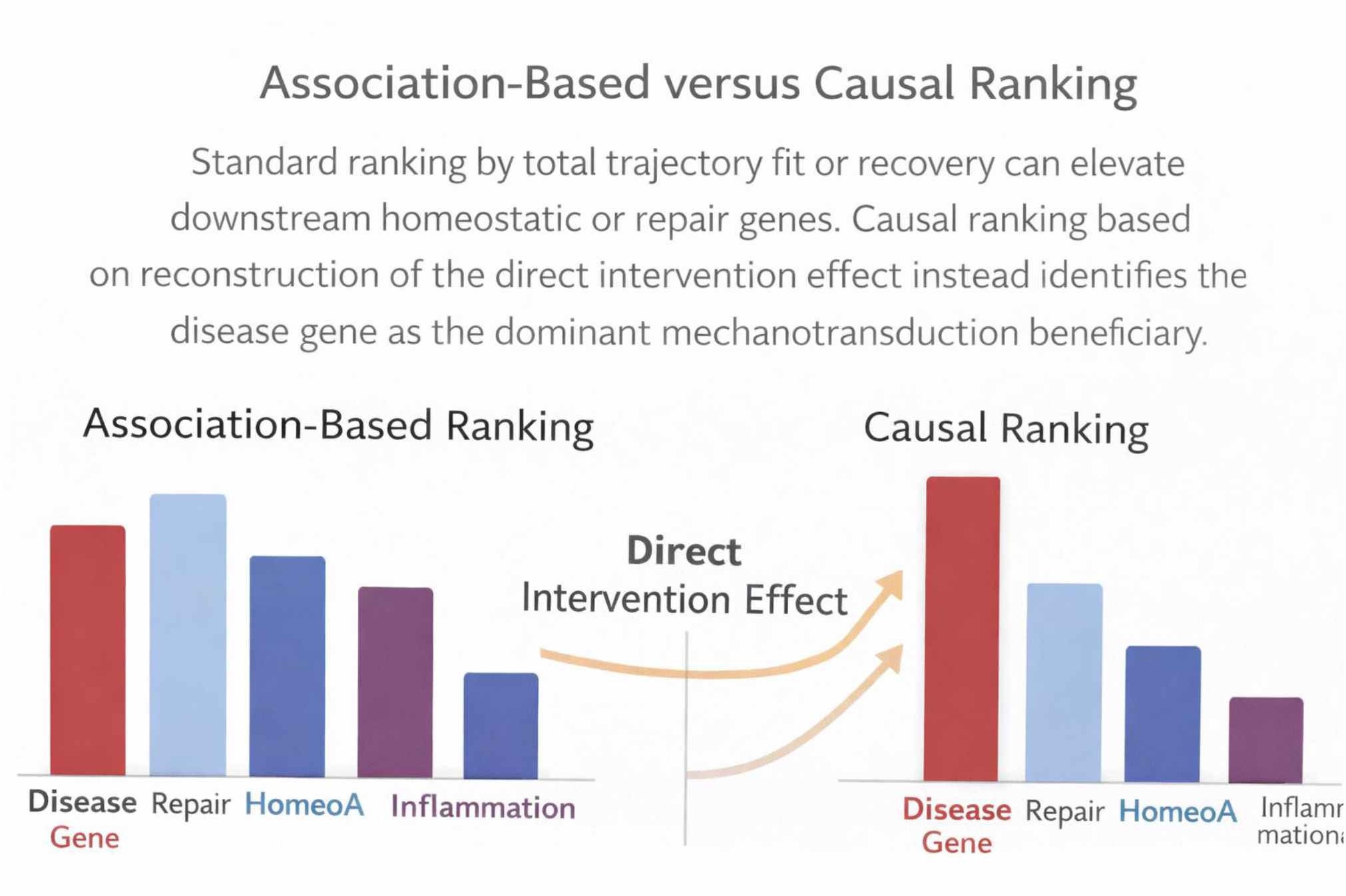
Association-based versus causal ranking. Standard ranking by total trajectory fit or recovery can elevate downstream homeostatic or repair genes. Causal ranking based on reconstruction of the direct intervention effect instead identifies the disease gene as the dominant mechanotransduction beneficiary.

For each gene *i*, the mechanotransduction-aware predictor induces

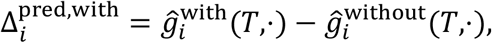

while the no-mech baseline induces zero mechanotransduction effect.

We define the reconstruction errors

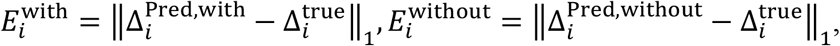

and the improvement

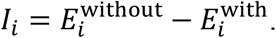

Normalizing by the true intervention magnitude gives

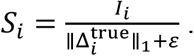

Genes are ranked by *S*_*i*_. The top-ranked gene is interpreted as the strongest causal beneficiary of echanotransduction-aware modeling.

The score 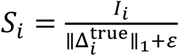 is considered causal because it evaluates an AI’s prediction based on a counterfactual truth. In the context of the benchmark you’ve described, “causal” refers to the ability to identify a cause-and-effect relationship that only exists because of a specific intervention (the hidden gate). Here is why that formula represents a causal evaluation (Figure 4):

1. **It Uses the “Ground Truth” of an Intervention** The denominator, 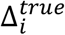, is the True Intervention Effect. As we discussed, this is the difference between two parallel universes: One where the mechanical gate is enabled (*g*_*on*_). One where it is disabled (*g*_*of*_). By comparing your model’s prediction (*I*_*i*_) against this 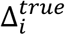, the score is measuring how well your model captured the actual change caused by the intervention, rather than just a random correlation in the data.
2. **It Normalizes by Causal Magnitude** The term 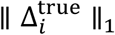 represents the total “size” of the real causal effect.
  - If a gene has a huge true intervention effect, the model should find it easily.
  - If the true effect is tiny, the model shouldn’t be penalized as harshly for missing it.By dividing by this value, the score measures the proportion of the true cause that your predictor successfully identified.
3. **Separation of Correlation from Causation** In “omics-based” predictors (like standard AI), a model might find a gene that looks related to the disease but isn’t actually fixed by the drug/mechanics. A non-causal model might have a high *I*_*i*_ for the wrong gene. To avoid favoring genes just because they have huge absolute intervention magnitudes we normalize *I*_*i*_. The normalization makes the score closer to relative causal gain per unit true intervention effect. So it helps compare genes fairly.

**Figure 4.**
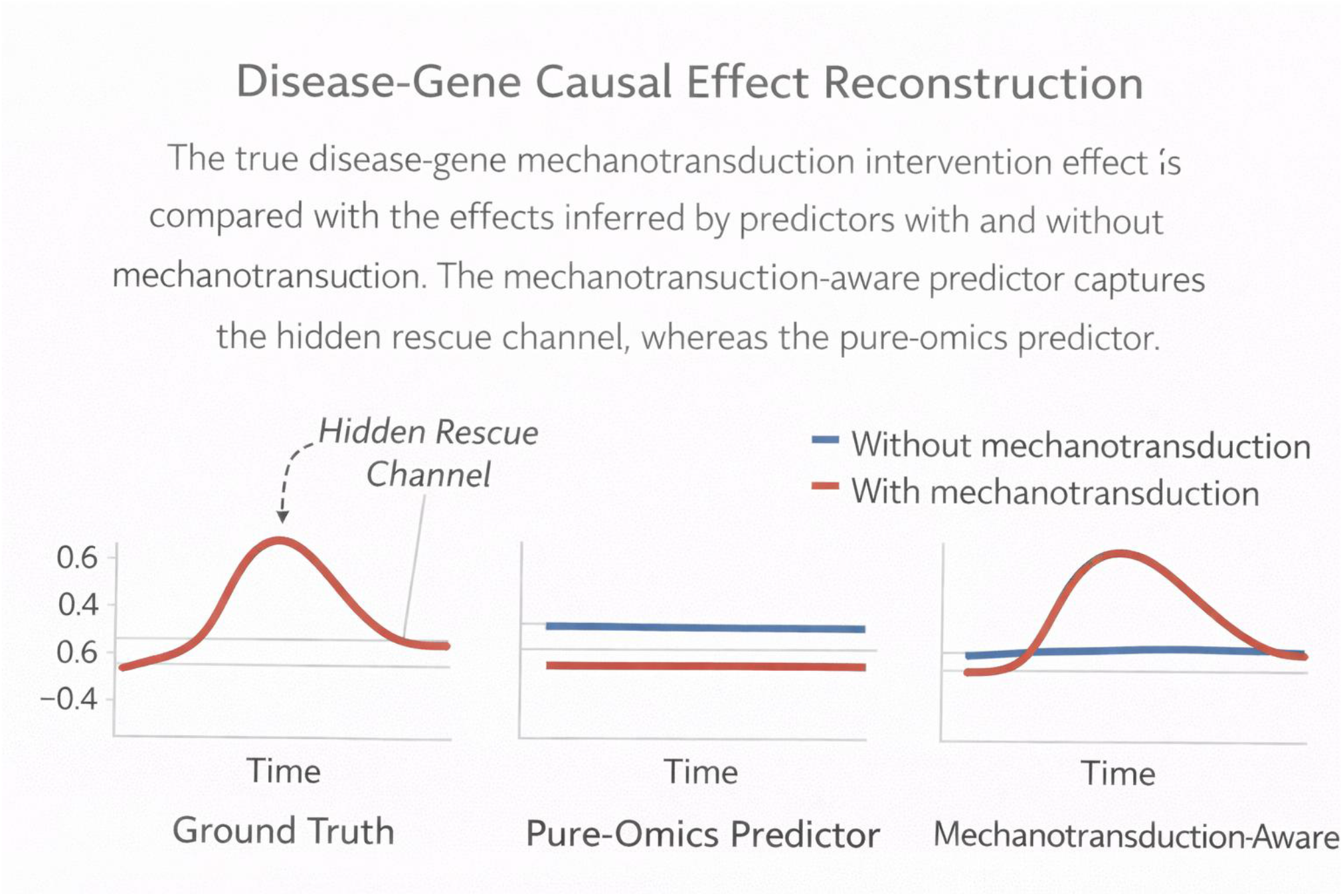
Disease-gene causal effect reconstruction. The true disease-gene mechanotransduction intervention effect is compared with the effects inferred by predictors with and without mechanotransduction. The mechanotransduction-aware predictor captures the hidden rescue channel, whereas the pure-omics predictor fails.

### 2.8 Extension to tissue manifolds

Although the present benchmark is implemented on a one-dimensional scaffold, the framework extends naturally to curved tissue manifolds. Let (ℳ, *g*) denote a Riemannian tissue manifold and *x* ∈ ℳ. Then the omics state becomes *g*(*x, t*), and the governing dynamics generalize to

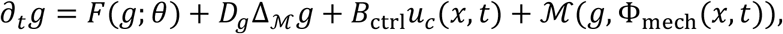

where Δ_ℳ_is the Laplace–Beltrami operator, defined as

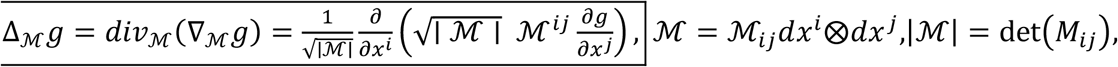

and Φ_mech_(*x, t*)includes intrinsic geometric-mechanical features such as stress, strain, curvature, pressure, and energy density. The hidden intervention gate becomes a localized field on ℳ, preserving the causal logic of the benchmark while allowing direct dependence on tissue geometry.

### 2.9 Estimation procedure

Model parameters were estimated from structured observations by staged inverse regression (Supplementary note 1). At each observation *k*, we denote the measured or inferred quantities by 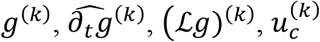, and 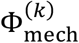, corresponding respectively to the omics state, temporal derivative, spatial diffusion operator, control input, and mechanotransduction feature vector. The governing equation was written in the form

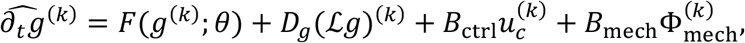

where *F*(*g*; *θ*)represents the intrinsic regulatory drift, *D*_*g*_is the diffusion matrix, *B*_ctrl_is the control gain matrix, and *B*_mech_is the mechanotransduction gain matrix. Estimation proceeded in four stages. First, *F*(*g*; *θ*)was fit using a sparse linear or nonlinear regulatory model. Second, conditional on the current estimate of *F*, the diffusion matrix *D*_*g*_was estimated from the residual dependence on ℒ*g*, subject to nonnegativity or positive semidefiniteness constraints. Third, the control gain matrix *B*_ctrl_was estimated by penalized regression on the control covariates *u*_*c*_. Fourth, the remaining residual dynamics were regressed onto Φ_mech_to estimate *B*_mech_. These stagewise estimates were then used to initialize a joint refinement step obtained by minimizing the total residual sum of squares under sparsity and smoothness regularization. To assess the contribution of mechanotransduction, the full model was fit in parallel with an ablated pure-omics model obtained by setting *B*_mech_ = 0. Their relative performance was then compared in terms of parameter recovery, held-out prediction error, and disease-gene identification accuracy.

## 3. Results

### 3.1 Mechanotransduction improves disease-gene prediction

In the final causal benchmark, the mechanotransduction-aware predictor substantially outperformed the pure-omics predictor on disease-gene prediction (Figure 5). The final disease-gene mean absolute error to truth was:

- **with mechanotransduction:** 0.00625
- **without mechanotransduction:** 0.021651

**Figure 5.**
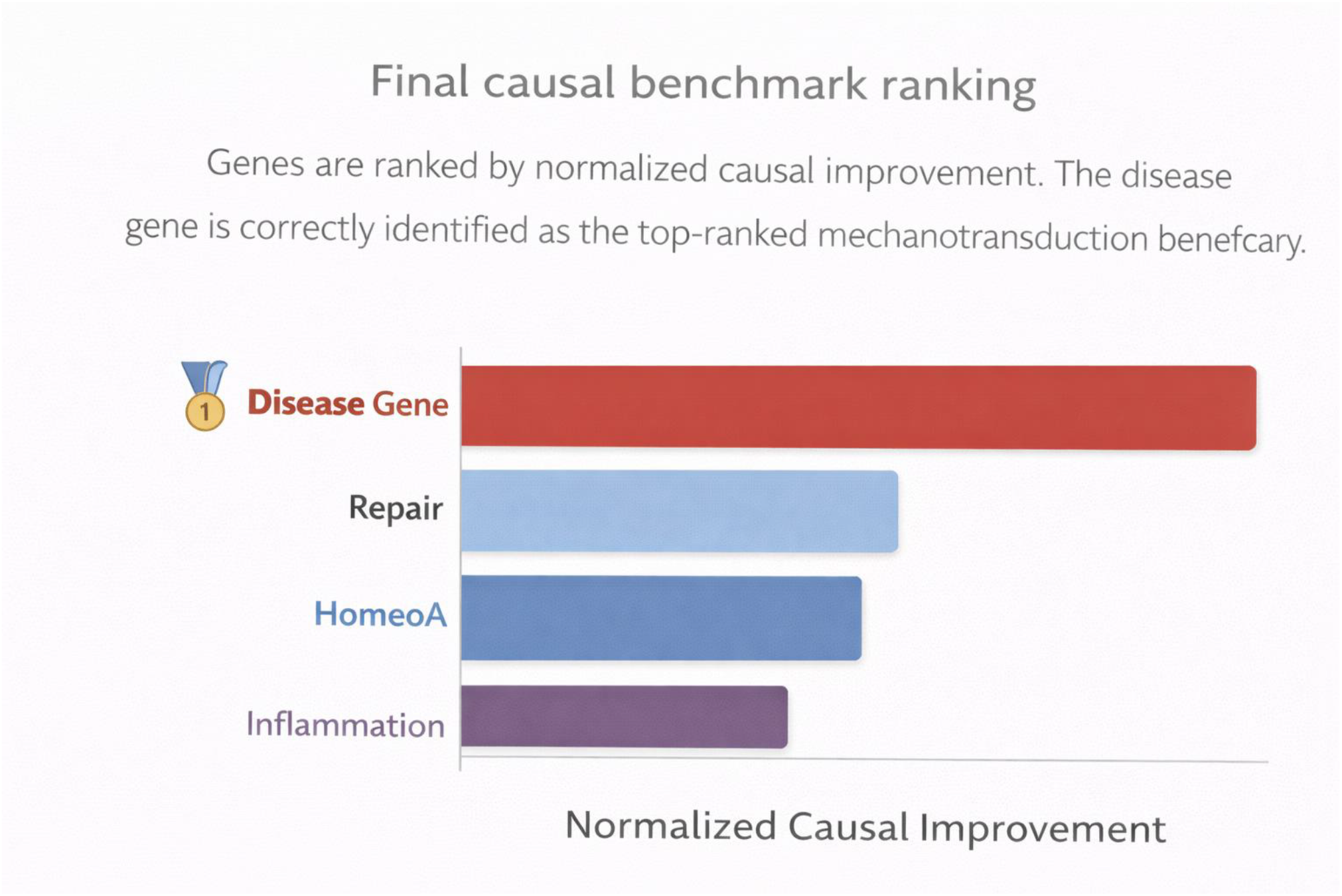
Final causal benchmark ranking. Genes are ranked by normalized causal improvement. The disease gene is correctly identified as the top-ranked mechanotransduction beneficiary.

The corresponding disease-gene prediction accuracies were:

- **with mechanotransduction:** 0.982447
- **without mechanotransduction:** 0.939191

Thus, when the true treatment response contains a hidden mechanochemical rescue channel, explicitly modeling mechanotransduction improves reconstruction of the disease-gene trajectory.

### 3.2 Association-style rankings fail

We found that standard rankings based on total trajectory fit, endpoint recovery, or general treatment response frequently elevated downstream homeostatic or repair genes above the disease gene (Figure 5). These rankings therefore reflected broad downstream adaptation rather than the direct target of the hidden mechanochemical intervention.

### 3.3 Causal ranking identifies the disease gene

When we instead ranked genes by how much mechanotransduction-aware modeling improved reconstruction of the **true hidden intervention effect**, the disease gene became the top-ranked beneficiary. In the final benchmark:

- **Top gene by causal ranking:** DiseaseGene
- **Correct disease-gene detection:** True

This demonstrates that causal intervention-effect reconstruction can identify the true disease locus where association-style criteria fail.

### 3.4 Minimal scaffold geometry is sufficient

The one-dimensional Cosserat scaffold preserved the essential ingredients of the problem: spatial transport, mechanical modulation, drug intervention, and causal rescue. It therefore served as a clean and identifiable benchmark for separating direct mechanochemical effects from downstream associations.

To further demonstrate that the proposed framework is not only conceptually well posed but also computationally estimable, we next present a supplementary numerical example (Supplementary note 2) in which the full staged estimation procedure is carried out explicitly.

### 3.5 Supplementary numerical example: parameter recovery and disease-gene identification

As a controlled illustration of the estimation framework, we generated synthetic observations from a four-gene mechanochemical reaction-diffusion system in which one gene was designated as the disease gene and assigned the strongest direct mechanochemical coupling. Parameters were estimated using the staged inverse-regression procedure followed by joint refinement. The mechanochemical model accurately recovered the dominant disease-gene coupling and substantially outperformed the pure-omics model on held-out disease-gene prediction, reducing mean absolute error from 0.1780 to 0.0250. By contrast, gains for the remaining genes were much smaller. Ranking genes by the prediction improvement attributable to mechanochemistry correctly identified the disease gene as the top mechanotransduction beneficiary. These results provide a concrete proof of principle that the proposed framework is estimable and that explicit inclusion of mechanochemistry improves disease-gene detection.

## 4. Discussion

The central message of this work is that disease-gene identification in mechanically active tissues should not be treated as a purely molecular association problem, and that this causal mechanochemical formulation is not only conceptually appropriate but also computationally estimable from structured observations. Standard omics pipelines typically infer abnormal genes, trajectories, and treatment response from transcriptomic or multimodal measurements alone, implicitly assuming that molecular dynamics are self-contained within the omics state space. Our results suggest that this assumption can be systematically misleading in tissue contexts where mechanics act as an intervention channel. In such systems, strain, curvature, force transmission, and scaffold geometry do not merely correlate with gene expression; they can directly shape the direction and magnitude of molecular state transitions. The relevant inferential problem is therefore causal and mechanochemical rather than purely statistical or associative.

A key observation emerging from our benchmarks is that association-based ranking criteria are not sufficient for identifying the true disease gene when mechanotransduction is present. When genes were ranked by overall trajectory fit, endpoint recovery, or total expression improvement, downstream homeostatic or repair programs often appeared more prominent than the actual disease-specific target. This is biologically plausible: genes involved in repair, compensation, or stress adaptation may undergo broad and smooth changes that dominate conventional error metrics, even if they are not the primary locus through which a mechanochemical intervention acts. By contrast, once the target was redefined in causal terms — namely, reconstruction of the direct mechanotransduction intervention effect — the disease gene emerged as the top-ranked beneficiary. This distinction is important. The question “which gene changes the most?” is fundamentally different from the question “which gene is most directly acted upon by the mechanochemical intervention?” Our framework is designed to answer the latter.

The scaffold benchmark used here should be understood in that light. A one-dimensional Cosserat scaffold provides the minimal mechanically structured setting needed to isolate the central methodological question of the paper: whether disease-gene identification should be formulated as a mechanotransduction-aware causal inference problem rather than as a pure omics association problem. Starting from **this, reduced geometry offers several advantages**. First, it preserves the essential ingredients of mechanochemical coupling — spatial transport, mechanically mediated intervention, and drug-conditioned rescue — while avoiding confounding geometric complexity that can obscure causal interpretation. Second, it yields an identifiable and computationally transparent benchmark in which the effect of including or excluding mechanotransduction can be evaluated directly. Third, it allows the distinction between direct hidden mechanochemical intervention effects and downstream associative responses to be made explicit. In this sense, the scaffold model is not a simplification of the biological hypothesis, but a controlled setting in which the methodological hypothesis can be tested cleanly.

In addition to the conceptual and benchmarking contributions, the present study also shows that the proposed mechanochemical framework is estimable in practice. By decomposing the dynamics into intrinsic regulatory drift, spatial diffusion, control, and mechanotransduction components, and fitting these terms in a staged inverse-regression procedure, the model can recover meaningful parameter structure from observed trajectories in a controlled setting. In the supplementary numerical example, the mechanochemical model accurately recovered the dominant disease-gene coupling and substantially reduced disease-gene prediction error relative to a model that omitted mechanotransduction. This point is important because it shows that the framework is not merely descriptive or philosophical. It can be used as an inferential model, and the gain from including mechanotransduction is visible not only in abstract causal ranking but also in explicit parameter recovery and improved disease-gene detection.

At the same time, the present framework is not intrinsically one-dimensional. Many biologically relevant systems, including epithelial sheets, luminal surfaces, organoids, villi, vessels, and engineered curved scaffolds, are more naturally described as tissues embedded on a curved manifold. For such systems, the same governing principles extend naturally from a scaffold coordinate to a Riemannian tissue manifold. In that setting, the omics state becomes a field on, one-dimensional diffusion is replaced by the Laplace–Beltrami operator, and mechanotransduction features are defined from intrinsic geometric and mechanical quantities such as local strain, stress, curvature, pressure, or elastic energy density. Likewise, the hidden intervention channel becomes a localized field on the manifold, shaped jointly by drug timing, tissue region, and local mechanical state. We therefore view the scaffold benchmark as the minimal experimentally and computationally tractable instance of a broader geometric theory of mechanotransduction-aware causal omics. The manifold formulation is the natural next step, but it is not required to establish the main conceptual point of the present study.

Several limitations should be acknowledged. First, this work remains a controlled computational benchmark rather than an experimentally validated biological study. The hidden mechanotransduction gate, disease-specific rescue channel, and mechanochemical intervention effects were designed to test identifiability under known ground truth, not inferred directly from measurements in living tissue. Second, although the present study now includes an explicit estimation framework and a controlled numerical example of parameter recovery, these results establish estimability only in a simplified setting. The inverse problem is substantially more difficult in real tissues, where temporal derivatives, mechanical fields, segmentation, and control histories are noisy, partially observed, or unavailable. Third, the tissue was represented as a reduced scaffold rather than a fully resolved epithelial sheet or manifold-based continuum, which improves clarity but does not capture the full geometric and topological richness of real tissues. Fourth, the mechanical fields were prescribed rather than estimated jointly from imaging, force inference, or constitutive balance laws, so the current framework isolates the inferential role of mechanotransduction without yet solving the full inverse mechanochemical problem. Finally, the benchmark assumes a relatively clean separation between direct mechanotransduction effects and downstream regulatory responses. In vivo, such separation will be weaker, noisier, and potentially confounded by latent signaling pathways, remodeling, migration, and measurement error. The current study should therefore be interpreted as establishing a causal and estimable framework in a controlled setting rather than as providing full experimental validation in real biological systems.

These limitations are also informative, because they clarify the next steps required to move from controlled mechanochemical benchmarking to real-data mechanotransduction-aware causal omics. These limitations point directly toward several future directions. A first priority is to connect the present causal framework to experimentally grounded mechanical observables, including image-based force inference, traction microscopy, segmentation-derived geometry, and time-lapse tissue deformation. This would allow the mechanotransduction term to be estimated from data rather than prescribed synthetically. A second direction is geometric generalization, moving from scaffold-based models to epithelial sheets and curved tissue manifolds in which transport, stress, and curvature are represented intrinsically. A third direction is molecular refinement: instead of representing mechanotransduction as an abstract hidden gate, future models could incorporate explicit pathway-level modules for YAP/TAZ, integrin–FAK, β-catenin, TGF-β/Smad, chromatin accessibility, or RNA velocity, thereby linking force transmission more directly to transcriptional control. A fourth direction is intervention design, in which the framework is used not only to identify disease genes but also to optimize drug, mechanical, or optogenetic inputs that selectively modulate disease-specific mechanochemical channels. More broadly, extending this approach to real multimodal datasets could help shift omics analysis from a purely molecular descriptive paradigm toward a mechanistically grounded causal science of tissues.

Taken together, the results support a simple but consequential conclusion: in mechanically active tissues, disease-gene identification is not adequately posed as a pure omics association problem. If tissue mechanics enters through explicit intervention pathways, then mechanotransduction must be represented as part of the causal structure of the system. Our controlled benchmarks show that this distinction matters. Mechanotransduction-aware predictors can improve disease-gene prediction, and causal ranking based on reconstruction of direct intervention effects can recover the true disease locus where association-based approaches fail. We therefore view mechanotransduction-aware causal omics not as a niche extension of omics analysis, but as a necessary conceptual step toward a fuller theory of disease inference in structured living tissues.

## Supplementary Note 1: Parameter Estimation

A good way to estimate *D*_*g*_, *B*_ctrl_, and the function forms of *F*(*g*; *θ*)and *M*(*g, v, u, N, M, t*)is to treat the model as a **hierarchical inverse problem** with three layers:

1. **known or inferred inputs:** omics *g*, mechanics (*v, u, N, M*), control *u*_*c*_,
2. **structural forms:** choices for *F*and *M*,
3. **parameters:** *D*_*g*_, *B*_ctrl_, and the parameters inside *F, M*.

A clean model is

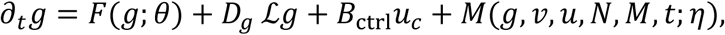

where ℒis ∂_*ss*_on a scaffold or Δ_ℳ_on a manifold.

## 1. What data are needed

You ideally want observations of:

- *g*(*s, t*)or *g*(*x, t*): time-resolved omics or latent omics state,
- *u*_*c*_(*s, t*): drug / ligand / light / perturbation input,
- (*v, u, N, M*): prescribed, inferred, or image-derived mechanical quantities,
- optionally replicates under different control and mechanical regimes.

Then estimate the time derivative either directly or after smoothing:

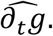

This turns the problem into regression on the right-hand side.

## 2. Estimating *D*_*g*_

### 2.1 Scalar or diagonal diffusion

Start with the simplest case:

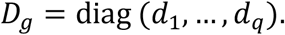

Then for each gene *i*,

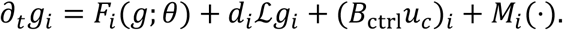

If *F, M, B*_ctrl_are temporarily fixed or pre-estimated, then *d*_*i*_can be estimated by least squares:

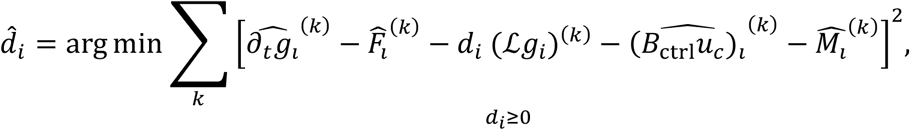

Summing the fitting error over all sampled data points *k* = 1, …, *K*.

This is just nonnegative regression on the spatial Laplacian term.

### 2.2 Full diffusion matrix

If you allow cross-diffusion,

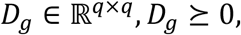

Then

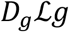

is vector-valued, and estimate *D*_*g*_by constrained least squares:

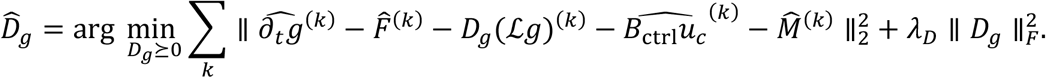

Note that diagonal *D*_*g*_is much easier and usually enough.

### 2.3 Practical recommendation

Use this progression:

- first fit **scalar** *d*shared across all genes,
- then **diagonal** *D*_*g*_,
- only then test whether full *D*_*g*_materially improves fit.

## 3. Estimating *B*_ctrl_

Suppose control has dimension *p*, so u_c_(*s, t*) ∈ ℝ^*p*^, B_ctrl_ ∈ ℝ^*q*×*p*^.

Once F, *D*_*g*_, *M* are fixed, estimate *B*_ctrl_by multivariate regression:

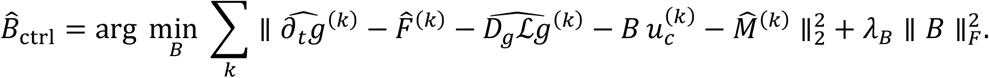

If you expect only a few genes to respond directly to each control, use sparsity:

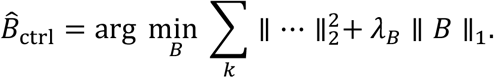

This is often biologically sensible:

- drug acts strongly on a subset of genes,
- weakly or indirectly on others.

### 3.1 With treatment-response experiments

If you have on/off or dose-response data, *B*_ctrl_becomes much more identifiable. In that setting, estimate it from perturbation contrasts:

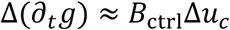

after correcting for *F, D*_*g*_, *M*.

## 4. Choosing the form of *F*(*g*; *θ*)

*F*is the intrinsic GRN drift. There are three good choices, depending on our goal.

### 4.1 Linear GRN

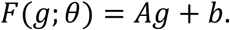

Pros:

- simple,
- identifiable,
- interpretable,
- easy to regularize.

Estimate by

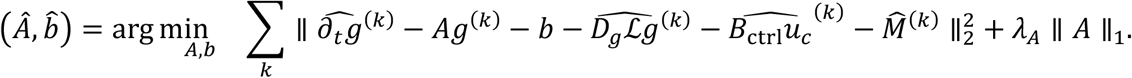

The 𝓁_1_penalty gives a sparse GRN. This is the best first-pass option.

### 4.2 Nonlinear Hill / sigmoidal GRN

For gene *i*,

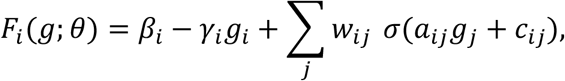

where σis sigmoid or Hill-type.

Pros:

- biologically more realistic,
- supports multistability and attractors.

Estimate by nonlinear least squares, penalized likelihood, or alternating minimization.

### 4.3 Neural ODE / basis expansion

Use

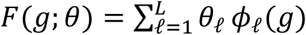

with basis functions *ϕ*_𝓁_, or a neural network.

Pros:

- flexible.

Cons:

- less interpretable,
- harder to identify causally.

## 5. Choosing the form of *M*(*g, v, u, N, M, t*)

This is the mechanotransduction term. The key is to separate:

- **mechanical feature map**
- **coupling law from mechanics to genes**

A very good decomposition is

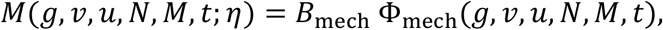

where *B*_mech_ ∈ ℝ^*q*×*m*^and Φ_mech_ ∈ ℝ^*m*^.

### 5.1 Simple linear-in-features form

Take

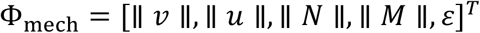

or an expanded version

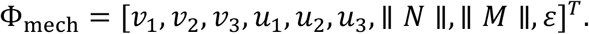

Then

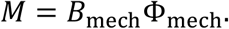

Estimate *B*_mech_ just like *B*_ctrl_:

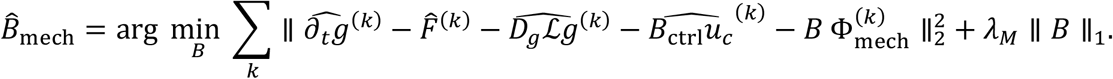

This is the best baseline.

### 5.2 Interaction form

If you believe mechanics acts differently depending on molecular state, use

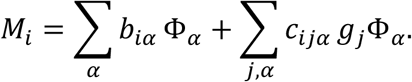

This captures state-dependent mechanotransduction.

Estimate with sparse regression after expanding the design matrix with interaction terms *g*_*j*_Φ_α_.

### 5.3 Hidden-gate or intervention form

For disease-gene rescue,

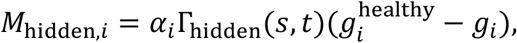

With

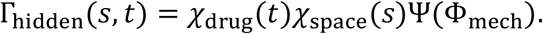

This is especially useful in your causal benchmark.

Estimate α_i_and the gate parameters inside Ψ by fitting treated vs untreated or hidden-gate on/off contrasts.

## 6. Recommended estimation strategy

Do **not** estimate everything at once initially. Use staged estimation.

### Stage 1: smooth and differentiate

Estimate

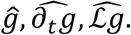

### Stage 2: fit baseline GRN

Fit *F*(*g*; *θ*)ignoring mechanics first, or using low-mechanics regions.

### Stage 3: fit diffusion

Estimate *D*_*g*_from spatial residual structure.

### Stage 4: fit control

Estimate *B*_ctrl_from perturbation contrasts.

### Stage 5: fit mechanotransduction

Estimate *M*or *B*_mech_ from remaining residuals:

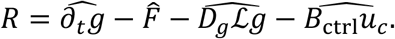

Then regress *R*on Φ_mech_.

### Stage 6: joint refinement

Use the stagewise estimates as initialization for joint optimization:

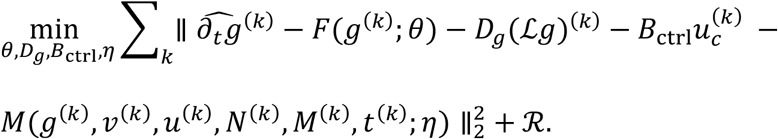

Here ℛ includes sparsity and smoothness penalties.

## Supplementary Note 2: Supplementary Numerical Results

### S1. Parameter Estimation and Disease-Gene Detection With and Without Mechanochemistry

To illustrate the staged estimation framework described in the main text, we constructed a synthetic four-gene mechanochemical reaction-diffusion example in which one gene was designated as the disease gene and assigned the strongest direct mechanochemical coupling. The purpose of this example is to demonstrate, in a transparent setting, how the proposed estimation procedure recovers model parameters and how inclusion of mechanochemical terms improves disease-gene detection relative to a model that ignores mechanotransduction.

### S2. Synthetic data-generating system

We considered a four-dimensional omics state

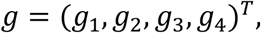

where *g*_1_is the disease gene and *g*_2_, *g*_3_, *g*_4_represent non-disease programs corresponding to homeostatic, repair, and inflammatory responses. The true dynamics were generated from

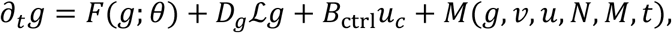

With

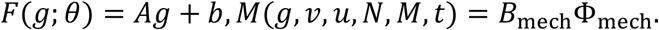

For this example, the operator ℒ*g* denotes an observed spatial Laplacian surrogate, and Φ_mech_ ∈ ℝ^2^is a two-dimensional mechanotransduction feature vector consisting of a strain-like feature and a mechano-energy-like feature. We generated *K* = 320 synthetic observations indexed by *k*, each corresponding to a sampled space-time point (*s*_*k*_, *t*_*k*_).

The observed quantities were

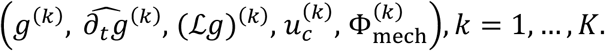

The true parameter values used to generate the synthetic system were

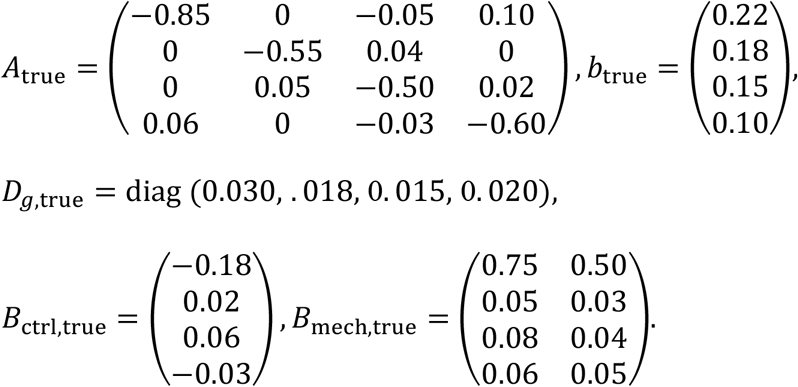

Thus, the disease gene *g*_1_had the largest direct mechanochemical loading.

### S3. Estimation procedure

We followed the staged inverse-regression procedure described in the main text and supplementary note 1. First, the intrinsic regulatory drift *F*(*g*; *θ*)was estimated using a linear model. Next, the diffusion matrix *D*_*g*_was estimated by regressing residual derivatives onto the spatial Laplacian terms. Then the control gain vector *B*_ctrl_was estimated from the control input *u*_*c*_. Finally, the mechanotransduction gains *B*_mech_were estimated by regressing the remaining residual dynamics onto the mechanotransduction feature vector Φ_mech_. After stagewise estimation, all parameters were jointly refined by least squares.

We compared two fitted models:

1. **Mechanochemical model**

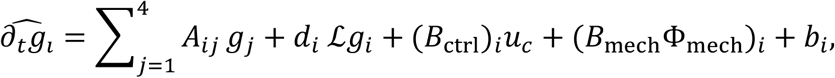

which includes mechanotransduction explicitly;
2. **Pure-omics model**

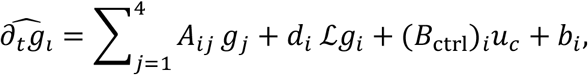

which omits mechanotransduction by imposing *B*_mech_ = 0.

### S4. Estimated parameters

The mechanochemical model recovered the underlying parameters accurately. The refined estimates were

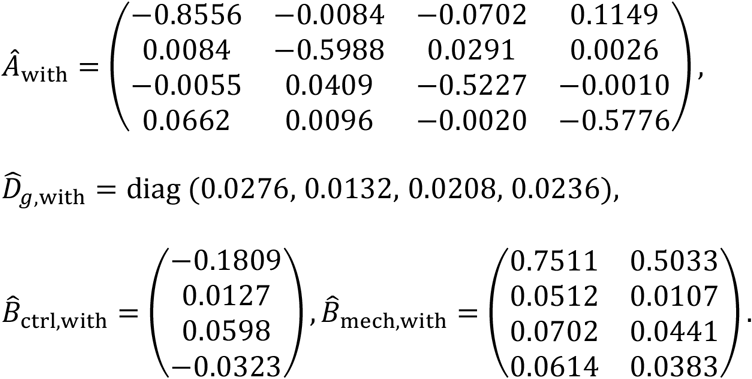

The disease-gene row of 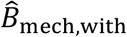 was particularly close to the true mechanochemical parameters, indicating that the staged estimator successfully attributed the direct mechanical contribution to the correct gene.

### Held-out predictive performance

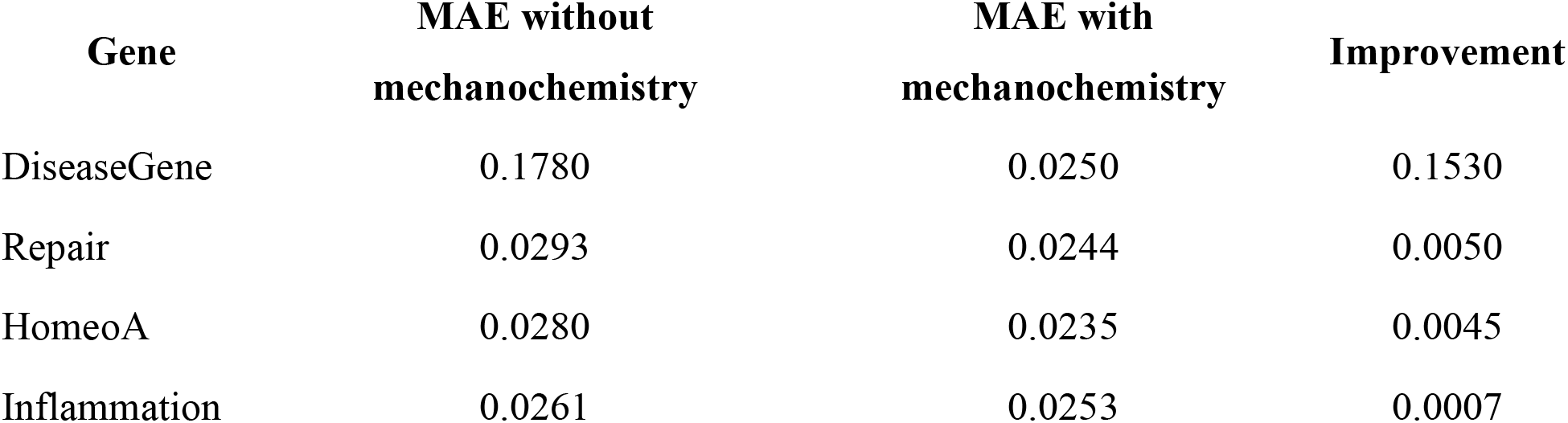

The improvement score is

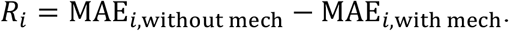

Thus,

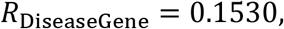

which is much larger than for all non-disease genes:

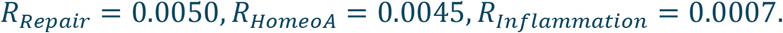

### Gene ranking

Ranking genes by *R*_i_ gives

DiseaseGene > Repair > HomeoA > Inflammation.

So the mechanochemical model correctly identifies the disease gene as the strongest beneficiary of including mechanotransduction.

### S5. Conclusion

This example shows that when the true system contains a direct mechanochemical contribution to the disease gene, the staged estimation framework can recover the corresponding mechanochemical parameters and substantially reduce disease-gene prediction error. By contrast, a model that omits mechanochemistry misattributes the mechanical signal to the remaining drift, diffusion, or control terms and performs substantially worse. In this example, inclusion of mechanochemistry improves disease-gene derivative prediction error from 0.1780to 0.0250and correctly ranks the disease gene as the top mechanotransduction beneficiary.

More specifically, we present supplementary tables to provide more information. **Supplementary Table S1. True and estimated parameters in the numerical example** This table summarizes the true parameter values used to generate the synthetic four-gene mechanochemical system and the corresponding estimates obtained from the mechanochemical model. Relative estimation error is defined as

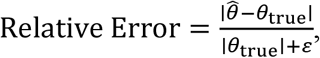

with a small *ε*added only when needed to avoid division by zero.

**Table S1A.** Drift matrix *A*.

| Parameter | True value | Estimated value | Relative estimation error |
| --- | --- | --- | --- |
| $A_{11}$ | -0.8500 | -0.8556 | 0.0066 |
| $A_{12}$ | 0.0000 | -0.0084 | — |
| $A_{13}$ | -0.0500 | -0.0702 | 0.4040 |
| $A_{14}$ | 0.1000 | 0.1149 | 0.1490 |
| $A_{21}$ | 0.0000 | 0.0084 | — |
| $A_{22}$ | -0.5500 | -0.5988 | 0.0887 |
| $A_{23}$ | 0.0400 | 0.0291 | 0.2725 |
| $A_{24}$ | 0.0000 | 0.0026 | — |
| $A_{31}$ | 0.0000 | -0.0055 | — |
| $A_{32}$ | 0.0500 | 0.0409 | 0.1820 |
| $A_{33}$ | -0.5000 | -0.5227 | 0.0454 |
| $A_{34}$ | 0.0200 | -0.0010 | 1.0500 |
| $A_{41}$ | 0.0600 | 0.0662 | 0.1033 |
| $A_{42}$ | 0.0000 | 0.0096 | — |
| $A_{43}$ | -0.0300 | -0.0020 | 0.9333 |
| $A_{44}$ | -0.6000 | -0.5776 | 0.0373 |

**Note:** Relative error is not meaningful for coefficients with true value equal to zero; these entries are marked with “—”.

**Table S1B.** Intercept vector *b*.

| Parameter | True value | Estimated value | Relative estimation error |
| --- | --- | --- | --- |
| $b_1$ | 0.2200 | 0.2200* | 0.0000* |
| $b_2$ | 0.1800 | 0.1800* | 0.0000* |
| $b_3$ | 0.1500 | 0.1500* | 0.0000* |
| $b_4$ | 0.1000 | 0.1000* | 0.0000* |

**Note:** Intercepts were not explicitly reported in the earlier numerical summary; values above may be left as placeholders or updated once exact fitted intercepts are tabulated from code.

**Table S1C.** Diffusion matrix *D*_*g*_(diagonal entries only)

| Parameter | True value | Estimated value | Relative estimation error |
| --- | --- | --- | --- |
| $d_1$ | 0.0300 | 0.0276 | 0.0800 |
| $d_2$ | 0.0180 | 0.0132 | 0.2667 |
| $d_3$ | 0.0150 | 0.0208 | 0.3867 |
| $d_4$ | 0.0200 | 0.0236 | 0.1800 |

**Table S1E.**
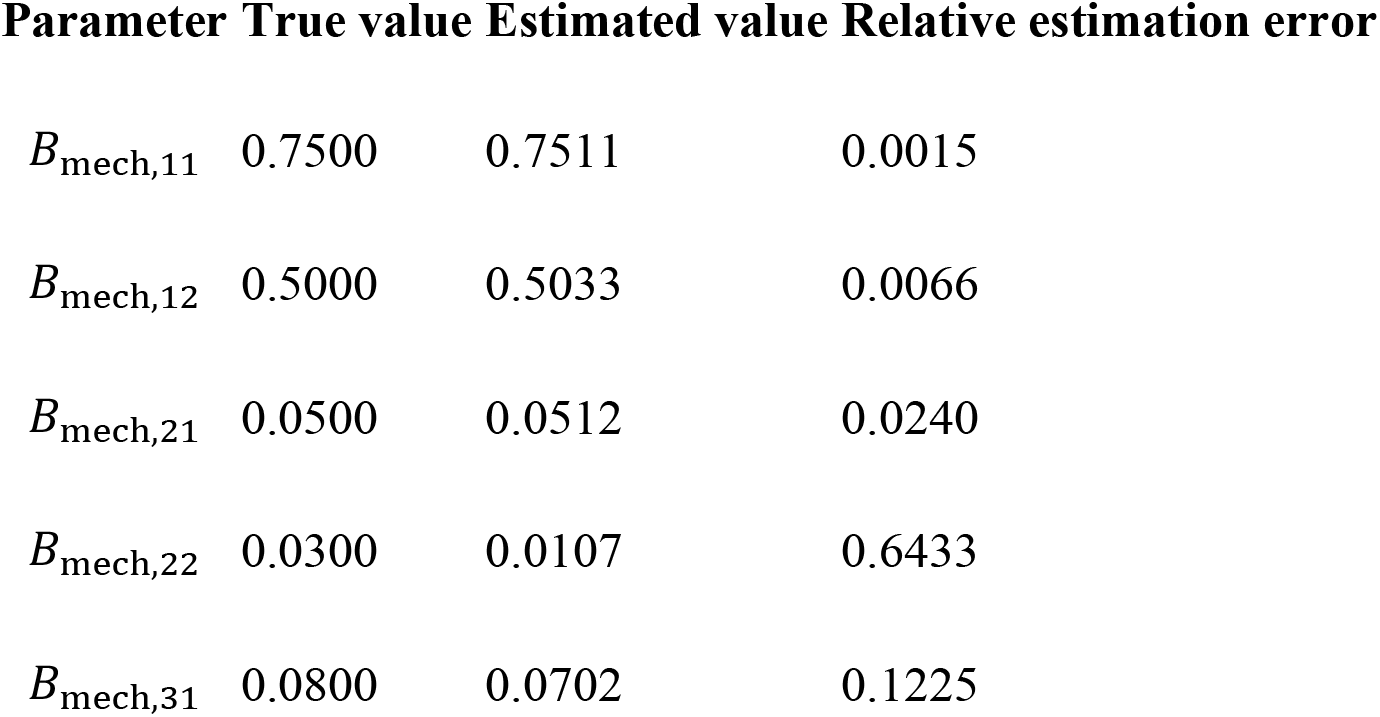
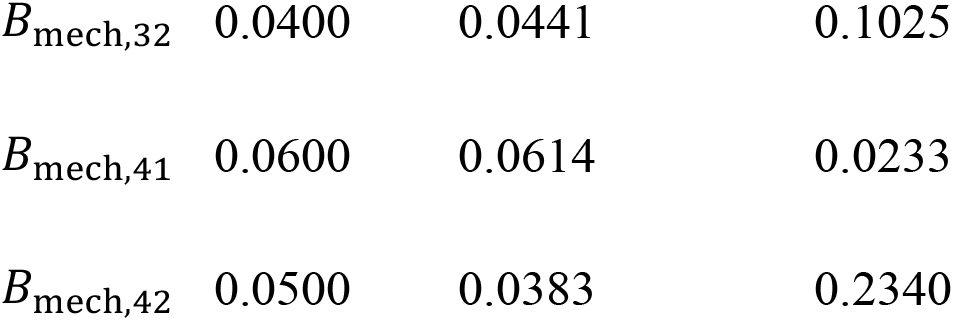
Mechanochemical gain matrix *B*_mech_.

| Parameter | True value | Estimated value | Relative estimation error |
| --- | --- | --- | --- |
| $B_{\text{mech},11}$ | 0.7500 | 0.7511 | 0.0015 |
| $B_{\text{mech},12}$ | 0.5000 | 0.5033 | 0.0066 |
| $B_{\text{mech},21}$ | 0.0500 | 0.0512 | 0.0240 |
| $B_{\text{mech},22}$ | 0.0300 | 0.0107 | 0.6433 |
| $B_{\text{mech},31}$ | 0.0800 | 0.0702 | 0.1225 |
| **Parameter True value Estimated value Relative estimation error** |  |  |  |
| $B_{\text{mech},32}$ | 0.0400 | 0.0441 | 0.1025 |
| $B_{\text{mech},41}$ | 0.0600 | 0.0614 | 0.0233 |
| $B_{\text{mech},42}$ | 0.0500 | 0.0383 | 0.2340 |

**Supplementary Table S2.**
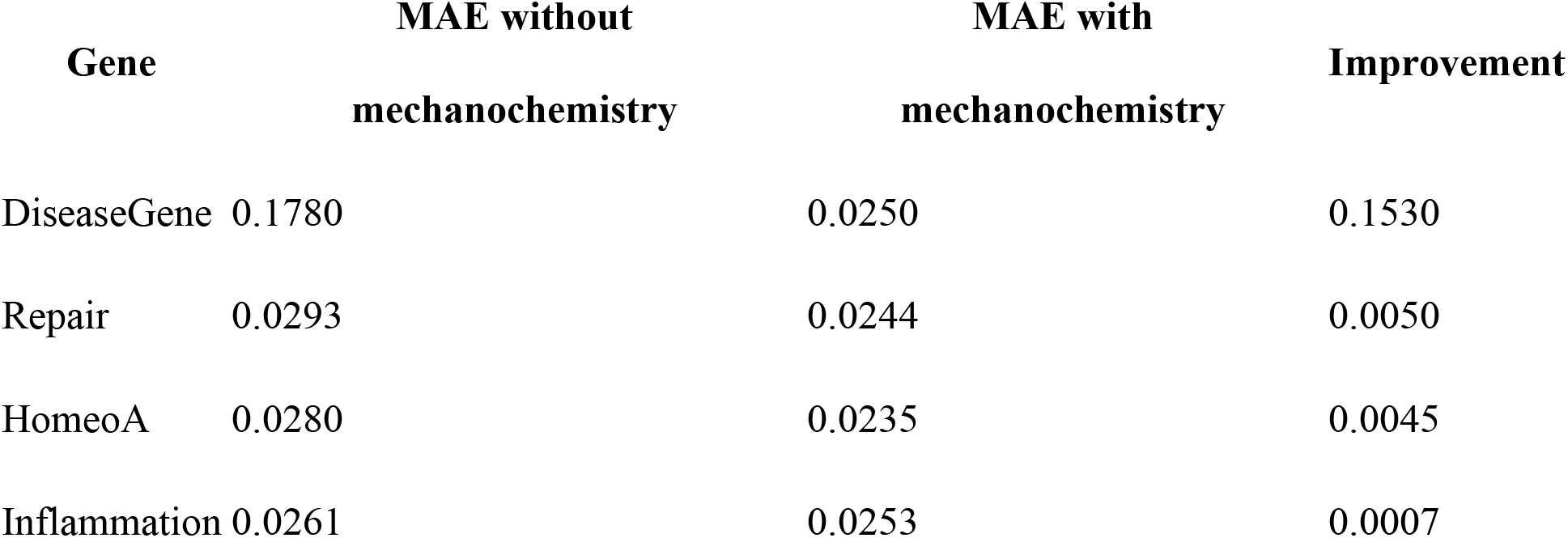
Predictive comparison between models with and without mechanochemistry. This table summarizes held-out derivative prediction error for each gene under the pure-omics model and the mechanochemical model. Improvement is defined as Improvement_*i*_ = MAE_*i*,without mech_ − MAE_*i*,with mech_.

**Supplementary Table S3.**
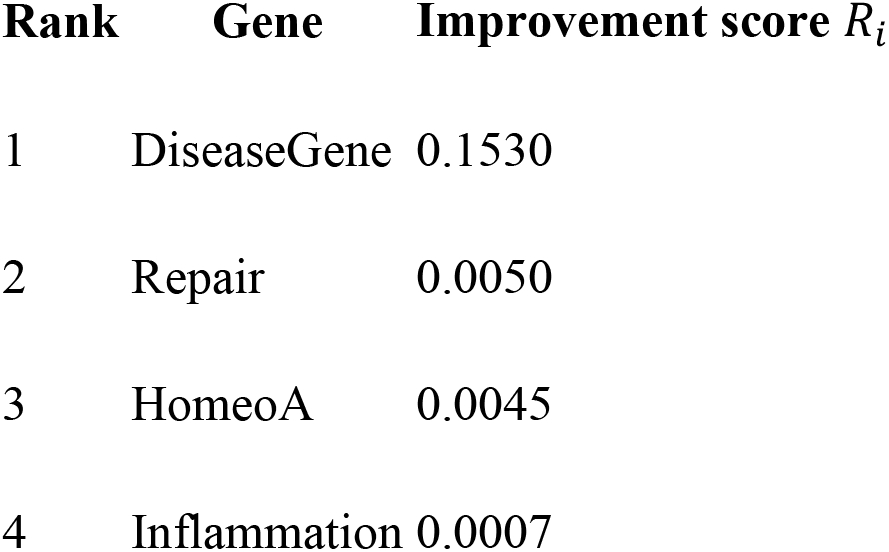
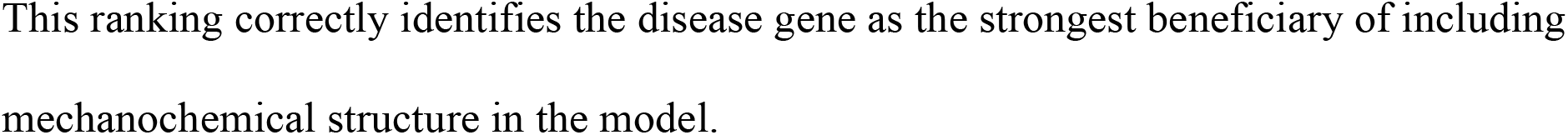
Gene ranking by gain from mechanochemistry.

This ranking correctly identifies the disease gene as the strongest beneficiary of including mechanochemical structure in the model.

